# miRBaseConverter: An R/Bioconductor Package for Converting and Retrieving miRNA Name, Accession, Sequence and Family Information in Different Versions of miRBase

**DOI:** 10.1101/407148

**Authors:** Taosheng Xu, Ning Su, Lin Liu, Junpeng Zhang, Hongqiang Wang, Weijia Zhang, Jie Gui, Kui Yu, Jiuyong Li, Thuc Duy Le

## Abstract

**Background:** miRBase is the primary repository for published miRNA sequence and annotation data, and serves as the “go-to” place for miRNA research. However, the definition and annotation of miRNAs have been changed significantly across different versions of miRBase. The changes cause inconsistency in miRNA related data between different databases and articles published at different times. Several tools have been developed for different purposes of querying and converting the information of miRNAs between different miRBase versions, but none of them individually can provide the comprehensive information about miRNAs in miRBase and users will need to use a number of different tools in their analyses.

**Results:** We introduce *miRBaseConverter,* an R package integrating the latest miRBase version 22 available in Bioconductor to provide a suite of functions for converting and retrieving miRNA name (ID), accession, sequence, species, version and family information in different versions of miRBase. The package is implemented in R and available under the GPL-2 license from the Bioconductor website (http://bioconductor.org/packages/miRBaseConverter/). A Shiny-based GUI suitable for non-R users is also available as a standalone application from the package and also as a web application at http://nugget.unisa.edu.au:3838/miRBaseConverter. *miRBaseConverter* has a built-in database for querying miRNA information in all species and for both pre-mature and mature miRNAs defined by miRBase. In addition, it is the first tool for batch querying the miRNA family information. The package aims to provide a comprehensive and easy-to-use tool for miRNA research community where researchers often utilize published miRNA data from different sources.

**Conclusions:** The Bioconductor package *miRBaseConverter* and the Shiny-based web application are presented to provide a suite of functions for converting and retrieving miRNA name, accession, sequence, species, version and family information in different versions of miRBase. The package will serve a wide range of applications in miRNA research and could provide a full view of the miRNAs of interest.

## Introduction

microRNAs (miRNAs) are short non-coding RNA molecules (about 22 nucleotides) that are widely encoded by eukaryotic nuclear DNA in plants and animals, and by viral DNA in some viruses [1, 2]. miRNAs are one of the vital and evolutionarily molecules that play an important role in post-transcriptional regulation by promoting degradation and repressing translation of their targets [3, 4]. As an important type of regulators, miRNAs are critical for gene expression that is associated with many biological and pathological processes. The amplification or genetic loss of some miRNAs can be the primary cause of many diseases, e.g. cancer, therefore making them valuable candidate biomarkers and receive lots of research attentions [5, 6, 7].

miRBase (http://www.mirbase.org/) is the home of miRNA data by providing a centralized system for assigning names and unique gene IDs for novel miRNAs [8, 9, 10]. It has become a primary repository of miRNA annotations and sequences for all species, a complete database of miRNA information, and the official rules for miRNA nomenclature. In addition, the miRNAs archived in miRBase are categorized into different miRNA families. The miRNA names and accessions confirmed by miRBase have been adopted in overwhelming of majority publications and miRNA-related molecular target databases since it was firstly established in 2002 [11, 12, 13].

miRBase has been constantly updated over the years, with 22 primary versions so far, due to the continuous study and deeper understanding of miRNAs [14, 15, 10, 16]. The latest miRBase version 22 was released in March 2018. In each new released miRBase version, the newly found miRNAs introduced in published literature were added in and the misannotated miRNAs in previous version were corrected or removed. To date, the miRNA numbers and the released time of the 22 primary miRBase versions are shown in Figure 1.

**Figure 1.**
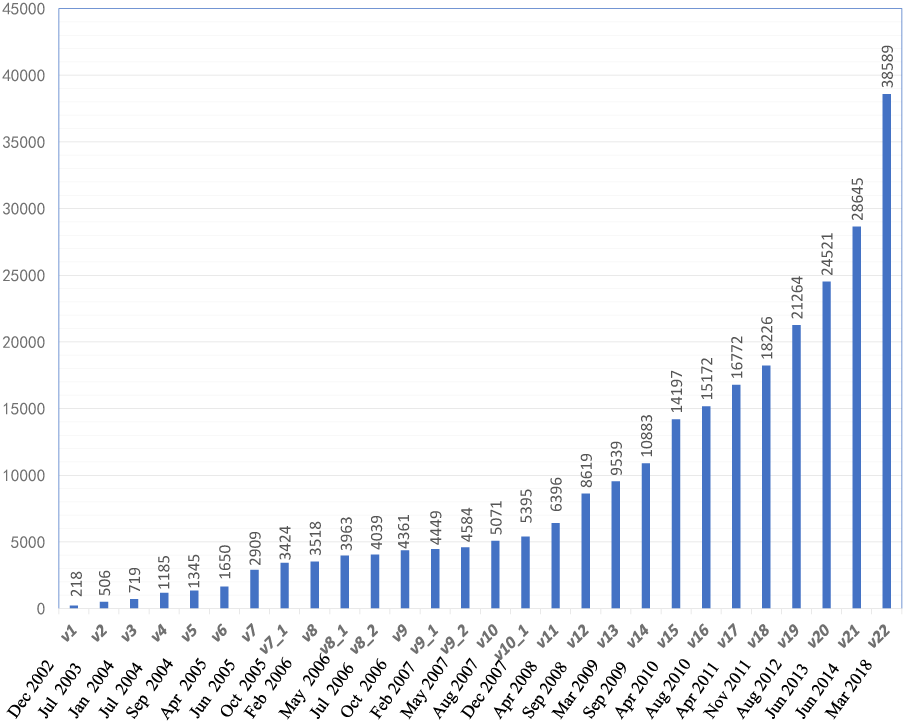
The number of hairpin precursor miRNAs accumulated in each of the miRBase versions. Then the hairpin precursor miRNAs could produce the two kinds of mature miRNAs from their 5’ and 3’ arms.

The definition and annotation of miRNAs vary significantly across different miRBase versions. However, corresponding changes have not been made in all other publicly available databases or publications. This issue of inconsistent miRNA annotations in public resources has become a barrier to the advancement of miRNA research. Researchers have to deal with inconsistent annotations of miRNAs in different resources, making it difficult for reusing previous research results for integrated analyses. As we know, the miRNA name annotation had a big change in miRBase versions 16,17,18 and 19 compared to the previous versions. For example, the “-as” nomenclature which had been designated as an antisense miRNA to another was abandoned in miRBase version 16. Meanwhile, the “miR*” designation was also gradually retired from this version. Then mature sequences from all Drosophila melanogaster precursors were designated as “-5p” and “-3p” in miRBase version 17. Mature sequences from all human, mouse and C. elegans precursors were firstly designated as “-5p” and “-3p” rather than “miR/miR*” in miRBase version 18. The mature miRNAs from all species have been designated as “-5p” and “-3p” instead of the mix “*”, “-5p” or “-3p” in miRBase version 19 and the distinction between “5p” and “3p” has been clearly defined since miRBase version 19.

There are several web-based or R-based tools for converting miRNA symbols between different versions of miRBase, including miRandola [17], miRSystem [18], miRiadne [19], miRLAB [20], miRNAmeConverter [21], miRBase Tracker [22], and miEAA [23]. Despite their usefulness, most of the tools are mainly focused on the miRNA name conversion between different miRBase versions to meet the most common research issues. Among them, miRNAmeConverter and miRBase Tracker offer more functions than the other existing tools. However, miRNAmeConverter is mainly designed for converting the annotations of mature miRNAs, and takes quite a long time to process the queries, while miRBase Tracker is a web application for accepting one miRNA retrieval each time. Hence, there is a lack of an easy-to-use, highly efficient, well-documented and comprehensive tool for retrieving miRNA information and converting the information in different versions of miRBase. To bridge this gap, we present the *miRBaseConverter R* package to provide a suite of functions for querying miRNA name, accession, sequence, species, version and family information in different versions of miRBase. Table 1 provides a summary of the features of *miRBaseConverter* and the other existing tools.

**Table 1.**
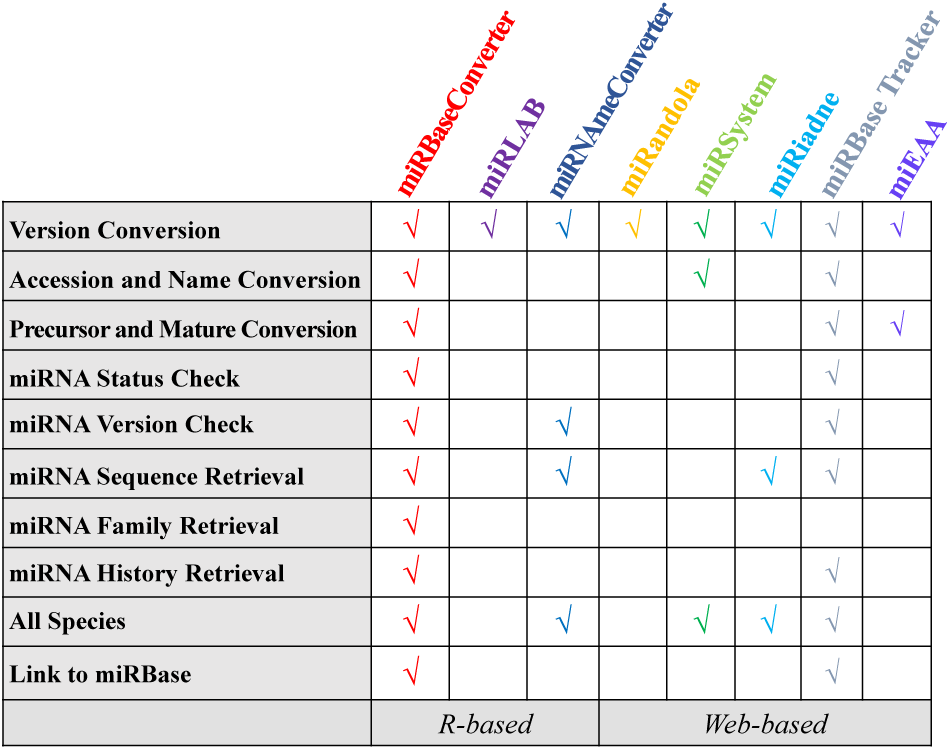
A summary of the different tools for information conversion among different miRBase versions

The *miRBaseConverter R* package is a full-scale tool for converting and retrieving information of all miRNAs defined in miRBase. Apart from the converting and retrieving functions, one of the outstanding features of *miRBaseConverter* is that it can process input containing the mix of premature and mature miRNA names, as well as accessions, in all species. The *miRBaseConverter R* package is easy to install without depending on other R packages. The aim of *miRBaseConverter* is to provide a handy and comprehensive R-based software tool and web application to the miRNA research community for integrating and analyzing miRNA datasets from different sources.

## MATERIALS AND METHODS

*miRBaseConverter* has a built-in database for converting and retrieving miRNA information in all species and for both pre-mature and mature miRNAs defined by miRBase. As shown in Figure 2, the *miRBaseConverter* R package provides four main types of functions: (1) miRNA check, (2) miRNA conversion, (3) miRNA information retrieval, and (4) miRBase online retrieval. In the following, we introduce the built-in database and the main functions of *miRBaseConverter* in detail.

**Figure 2.**
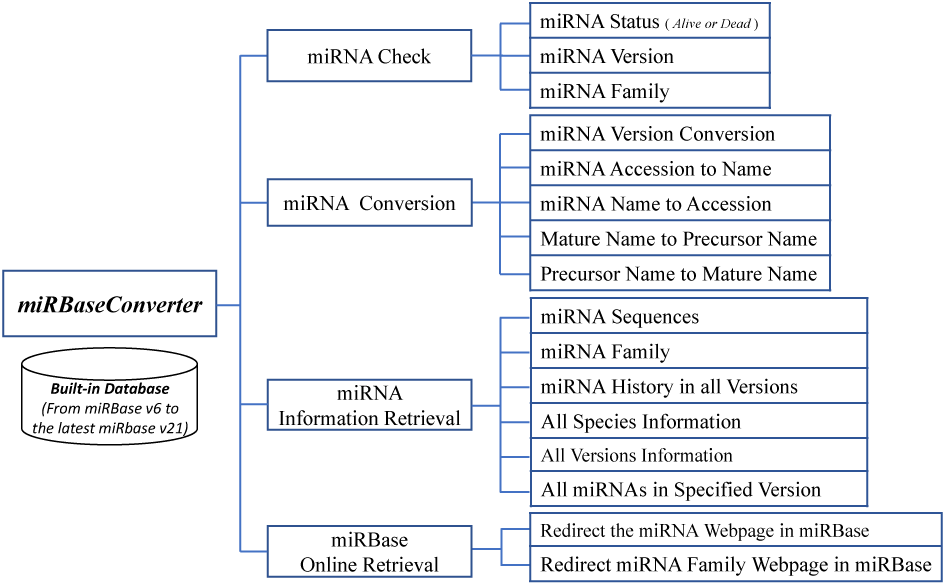
The functional framework of *miRBaseConverter*.

### The Built-in Database

We have constructed a built-in database for *miRBaseConverter* to include the miRNA information from miRBase version 6 to version 22 (the latest version). The miRNA names, accessions, sequences, families, species and other information in the different versions of miRBase are organized and stored in an index structure. Each of the same types of miRNA data (e.g. names, accessions, or sequences) is summarised together and each entry is assigned a unique index. Then the mapping tables between different data types are generated based on the indexes of entries. The stored information in the database can be greatly reduced by only keeping one copy of the same information in different versions of miRBase. Finally, the database is compressed to have a very small size (∼ 1.69 MB), making it easy to be embedded into the R package and be loaded to R workspace in runtime. The index structure of the database greatly improves the retrieval efficiency for a large amount of miRNAs. The built-in database is synchronized regularly with the miRBase repository.

### miRNA Check

*miRBaseConverter* can be applied to fast check the miRNA status (Alive or Dead), the most probable miRBase version number, and the family information of a list of miRNAs. Due to the limitations at the time when a miRBase version was created, there were some misannotated and duplicated sequences included in different versions of miRBase. These misdefined miRNAs were always killed in the following updated miRBase versions. However, some databases or research works may still include the results related to these misdefined miRNAs. There is a demand for addressing this issue to rapidly screen dead miRNAs. The version number for a list of miRNAs is the valuable knowledge for downstream analysis. *miRBaseConverter* provides the function to check the most probable miRBase version for the input miRNA list and gives the matched proportions in all miRBase versions. Meanwhile, *miRBaseConverter* is the first tool for batch querying miRNA family information, which is important for functional enrichment research of miRNAs. The defined functions in *miRBaseConverter* for miRNA check are *checkMiRNAAlive(), checkMiRNAVersion()* and *checkMiRNAFamily()*.

### miRNA Conversion

*miRBaseConverter* provides five kinds of miRNA conversion functions for all species: miRNA name to accession, accession to name, name to name in different versions, precursor name to mature name and mature name to precursor name. The corresponding functions are *miRNAJNameToAccession(), miRNA jiccessionToName(), miRNAVersionConvert(), miRNA_PrecursorToMature()* and *miRNA JMatureToPrecursor().* The fast batch conversion between miRNA name and accession is the basic need for miRNA research. An accession is the unique and fixed identifier assigned to a sequence record of a miRNA and therefore miRNA accessions are adopted in most miRNA-related databases. However, miRNA names have better readability than accessions for researchers in most cases. Meanwhile, there is a great demand for the conversion of miRNA names in different versions. For example, Tarbase v7 [24] and miRTarbase v7 [25] use miRBase version 18 and version 21, respectively. When we integrate those resources into our analyses, we need to convert the miRNA names in the two databases to the names used in the same version of miRBase. In addition, the conversion between precursor names and mature names is required in some cases, e.g. when integrating data from TCGA and GEO into the same analysis. *miRBaseConverter* offers a wide range of conversion tools. A number of examples illustrating the conversions are provided in the manual of the package.

### miRNA Information Retrieval

We implement functions for the retrieval of all kinds of miRNA information, including the basic statistical information of all miRBase versions, the species information involved in all miRBase versions, the detailed history information of a miRNA in all miRBase versions, the family information of miRNAs, the sequence information of miRNAs and the complete data table of all miRNAs in a specified miRBase version. The defined functions for miRNA information retrieval are *getAllMiRNAs(), getAllSpecies(), getAllVersionInfo(), getMiRNAHistory(), getMiRNASequence(),* and *getMiRNATable().* By calling the functions implemented in *miRBaseConverter* to retrieve the information of a miRNA or a list of miRNAs, it is easy to get the particularly detailed knowledge of the miRNAs of interests.

### miRBase Online Retrieval

*miRBaseConverter* provides two functions *goTojniRBase()* and *goTo-miRNAFamily()* to redirect users to miRBase web server for retrieving more detailed information (miRNA webpages and miRNA family webpages) of miRNAs. The operation of miRNA online information retrieval gives a comprehensive perspective of the miRNAs of interest and greatly helps with understanding the miRNAs.

## RESULTS

The *miRBaseConverter* package implemented in R is designed for the miRNA research community to assist with converting and retrieving miRNA information in different versions of miRBase. *miRBaseConverter* provides a suite of functions for querying miRNA name, accession, sequence, species, version and family information. The package is released under under the General Public License (GPLv2) and freely available from Bioconductor (3.5) repository [26]. In addition, we also provide the implemented functions and the characteristics via the online web application. In the following, we present the basic usage of *miRBaseConverter*, the shiny-based web application and two typical case studies to demonstrate the applications of *miRBaseConverter* with different purposes. The relevant datasets for the case studies are provided as supplementary data and available at https://taoshengxu.github.io/ExperimentData/miRBaseConverter.

### Using *miRBaseConverter* in R environment

The user manual of *miRBaseConverter* provides a detailed document to illustrate how to use each of the functions of the package, which can be accessed in the Bioconductor repository. Here we demonstrate a use case of the *miRBaseConverter* R package for the version upgrade of a list of miRNAs. The original miRBase version of the candidate miRNAs is always missing or is not provided in most actual cases. The function *checkMiRNAVersion()* can be applied firstly to check the most probable miRBase version number for the list of miRNAs. Then we use the miRNA accession (as the transforming bridge) to exactly match each of the miRNAs which might have been defined in different miRBase versions. The following are the code chunks and the output for converting a list of miRNAs to the latest miRBase version 22. The detailed user manual can be accessed via Bioconductor website (https://bioconductor.org/packages/release/bioc/manuals/miRBaseConverter/man/miRBaseConverter.pdf).

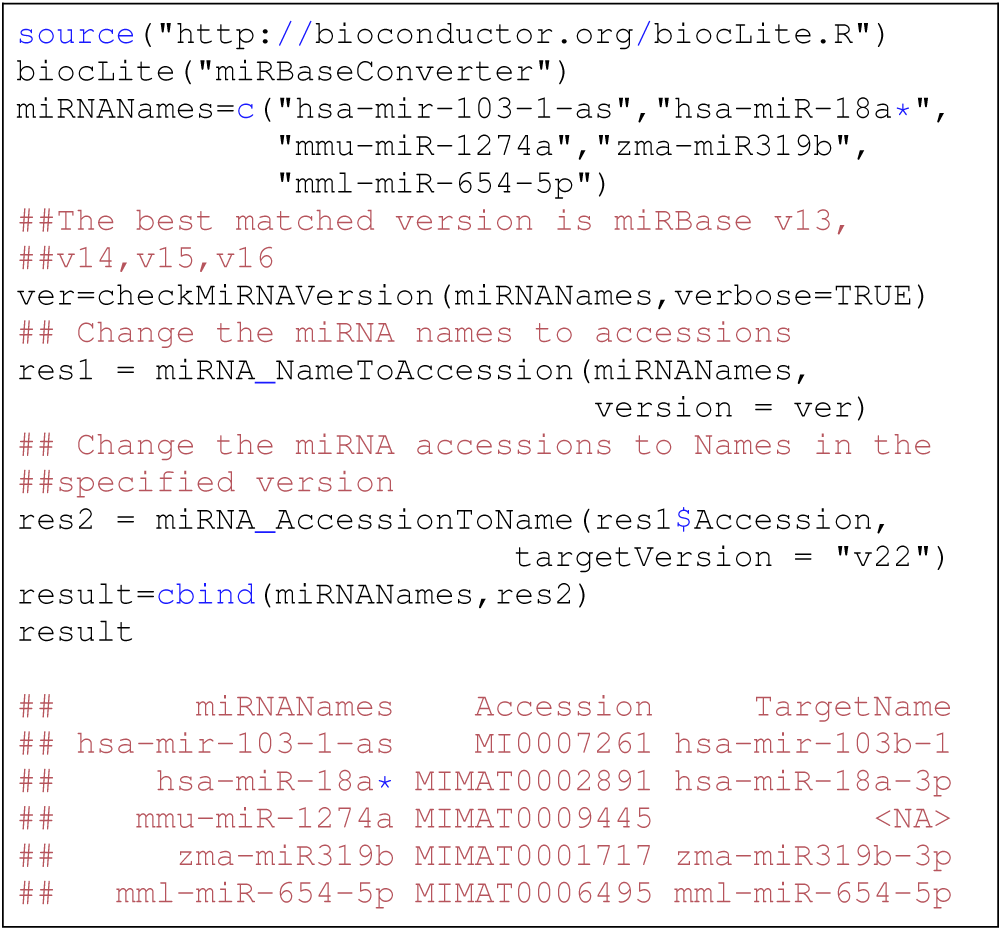

The above example shows that *miRBaseConverter* can successfully process the input precursor and mature miRNAs from different species at the same time. For example, the precursor miRNA is *hsa-mir-103-1-as* in miRBase version 16 converted to *hsa-mir-103b-1* in miRBase version 22, and the mature miRNA *mmu-miR-1274a* in miRBase version 16 is removed in miRBase version 22. More examples for miRNA information retrieval and version conversion is presented in the vignette of *miRBaseConverter*, which is available in Additional file 1.

### Retrieving and converting the information of a list of miRNAs in Shiny-based web application

We have designed a web application with a GUI to provide non-R users a convenient way to retrieve and convert miRNA information in different versions of miRBase without depending on R runtime environment. The Shiny-based web application provides a shell for the *miRBaseConverter* package which runs in web browsers. The GUI adopts a left-right structure with a concise performance as shown in Fig 3.

**Figure 3.**
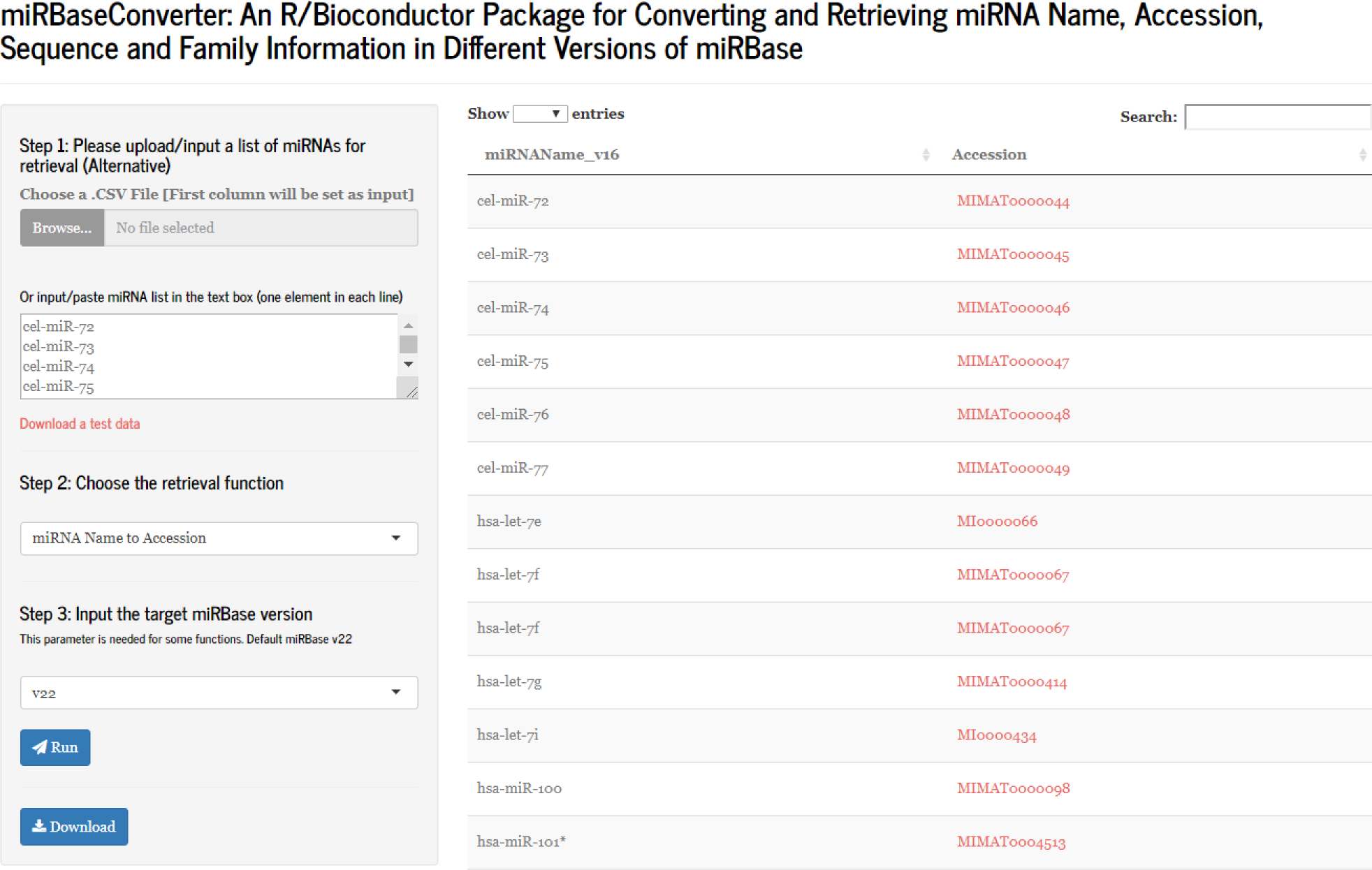
The GUI of the Shiny-based web application for *miRBaseConverter* package.

There are three main steps for the retrieval of a list of miRNAs as shown in the left pane of the GUI in Fig 3. Users are asked to upload or input the miRNA list (miRNA names or miRNA accessions) to be processed firstly. Then there is a drop-down box providing a list of main functions for the miRNA retrieval and conversion in the second step, which covers most functions in the *miRBaseConverter* R package. In the third step, a additional parameter which is the target miRBase version is needed for some of the main functions listed in the second step. After finalising all the three steps, the “Run” button is used for submitting the input content to server for result retrieval.

On the right-hand side of the GUI, there is a result table presenting the retrieval and conversion information of the input miRNA list. The records in the result table strictly correspond to the order of the miRNAs in the input list. The miRNA accessions or family accessions in the result table are implanted hyperlinks to easily redirect users to the miRBase web server for the detailed information about the miRNAs of interest. In addition, the web application provides the function to export the miRNA result table (a .csv file) for further study. The functions in the *miRBaseConverter* R package are called when users submit the retrieving or converting request from the web GUI, so the retrieval and conversion via the web application also have the quick response as using *miRBaseConverter* in the R environment.

### Case study 1: Constructing gene regulatory network of mRNAs, TFs and miRNAs

In this scenario, we present the use of *miRBaseConverter* for constructing a gene regulatory network including mRNAs, TFs and miRNAs based on a variety of interaction databases. There are quite a number of experimentally confirmed and putative databases that reveal the regulatory relationships among biological molecules. Information from these databases can be integrated together to form a comprehensive biological regulatory network. However, the miRNA names in different databases are inconsistent as they use different versions of miRBase for the miRNA annotation. Therefore, a unified miRNA annotation is the foundation for constructing the gene regulatory network. *miRBaseConverter* offers a handy tool in high throughput workflow to unify the miRNA names for various miRNA interaction databases. Here we choose some representative databases used for each type of interactions in our analysis. The detailed information of the interaction databases is summarised in Table 2.

**Table 2.**
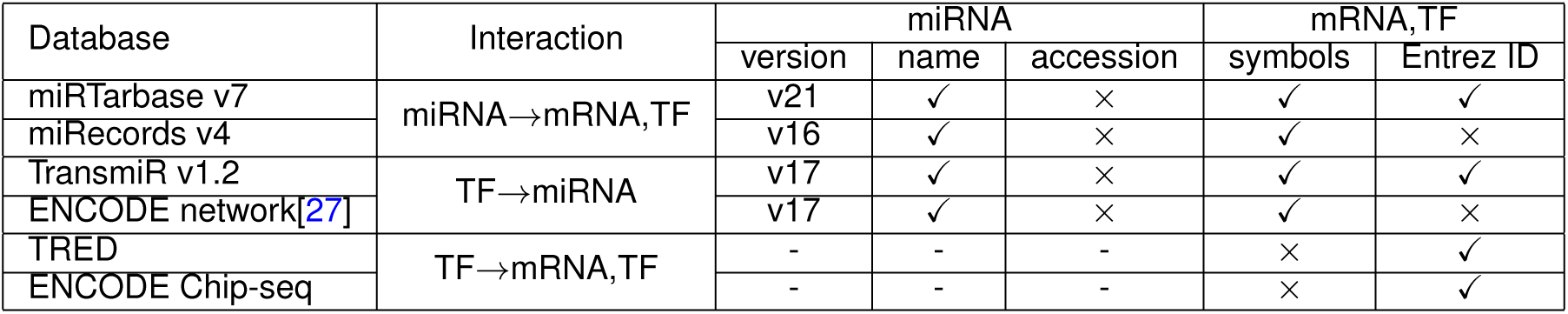
The main features of the interaction databases for constructing gene regulatory network

| Database | Interaction | miRNA |  |  | mRNA,TF |  |
| --- | --- | --- | --- | --- | --- | --- |
|  |  | version | name | accession | symbols | Entrez ID |
| miRTarbase v7 | miRNA→mRNA,TF | v21 | ✓ | × | ✓ | ✓ |
| miRecords v4 |  | v16 | ✓ | × | ✓ | × |
| TransmiR v1.2 | TF→miRNA | v17 | ✓ | × | ✓ | ✓ |
| ENCODE network[27] |  | v17 | ✓ | × | ✓ | × |
| TRED | TF→mRNA,TF | - | - | - | × | ✓ |
| ENCODE Chip-seq |  | - | - | - | × | ✓ |

miRTarbase is an experimentally validated and widely used miRNA-target interaction database. miRTarbase v7 collects 502652 records of human miRNA-mRNA interaction pairs, where miRNA names and the gene symbol and Entrez ID information of the target mRNAs are provided [25]. But it doesn’t involve the miRNA accession. As mentioned above, a miRNA accession is the identifier that defines a miRNA uniquely in miRBase. Therefore, it is more reliable to use miRNA accessions when using different miRNA databases at the same time. This kind of batch conversion is the basic function provided by *miRBaseConverter*. miRecords is a resource for animal miRNA-target interactions [28]. We use the validated targets in miRecords for our experiments in this case study. The data of miRecords v4 includes the miRNA names and the target gene symbols. The miRNA accessions and gene Entrez IDs are missing. We apply *miRBaseConverter* and the *org.Hs.eg.db* package to retrieve the miRNA accessions and gene Entrez IDs, respectively. TransmiR is a database of TF-miRNA regulations [29]. The TF genes in TransmiR v1.2 are with the Entrez ID information and the miRNAs information is provided as miRNA precursor names. However, mature miRNAs are commonly adopted as the regulatory factors in most interaction databases, such as miR-Tarbase v7 and miRecords v4 introduced previously. As the many studies about gene regulatory network, we also only apply mature miRNAs as the regulatory factors in the gene regulatory network to be constructed. Therefore, the miRNA precursors in TransmiR v1.2 need to be converted to the mature miRNAs. *miRBaseConverter* is an appropriate tool for the conversion of miRNA precursor names and mature names. The ENCODE research work [27] provides a highly credible collection of regulatory interactions between TFs and miRNAs in its supplement material. The interaction data can be downloaded from http://encodenets.gersteinlab.org/. Similar to miRecords, we apply *miRBaseConverter* and the *org.Hs.eg.db* package to retrieve the miRNA accessions and Entrez IDs for the interaction data, respectively. We get the information in Transcriptional Regulatory Element Database (TRED) [30] and ENCODE Chip-seq data from a collection of sources (https://github.com/slowkow/tftargets).

Due to the fact that miRTarbase v7 adopted the relatively new miRBase version (v21) than the other miRNA-related databases, we unify the miRNAs in all the miRNA interaction databases to miRBase v21. Firstly, the large number of miRNA names in miRTarbase v7 can be converted to the corresponding miRNA accessions using the function *miRNA_NameToAccession()* in *miRBaseConverter.* Before the conversion, we apply the *check-MiRNAVersion()* function to have a double-check on the miRBase version of the miRNA name list to be processed. Then the functions *miRNA JNameToAccession()* and *miRNA JAccessionToName()* are used for converting the miRNAs involved in miRecords v4 (miRBase v16) to the names and accessions in miRBase v21. For the miRNAs in other databases, we process them in the similar way to have a consistent miRBase version for downstream integration analysis.

By applying *miRBaseConverter* and other gene annotation R packages, all the miRNAs and genes involved in the interaction databases have been adopted their own unified annotation, respectively. Therefore, the regulatory interactions from all the databases are combined together to form a two-column data table representing the complex gene regulatory network. The two-column data table structure is the most common representation of a gene regulatory network. A directed graph can be generated to visualize the gene regulatory network. The nodes are the biological features (miRNAs, TFs and mRNAs) and the edges are the regulatory interactions between the features. For illustration, in Fig 4, we show the TP53 gene sub-network extracted from the constructed complex gene regulatory network by using Cytoscape [31]. The TP53 gene encodes the tumor suppressor protein p53 that regulates the cell cycle to prevent cancer formation [32]. The p53 pathway and its interaction networks are promising sources for cancer research [33, 34]. We further generate a sparse matrix as the mathematical representation for the directed graph. The sparse matrix can be applied as the input of most computational biology algorithms, such as GeneRank [35], NCIS [36] and WSNF [37]. The complete processing scripts and datasets for this case study is provided in the online supplementary data.

**Figure 4.**
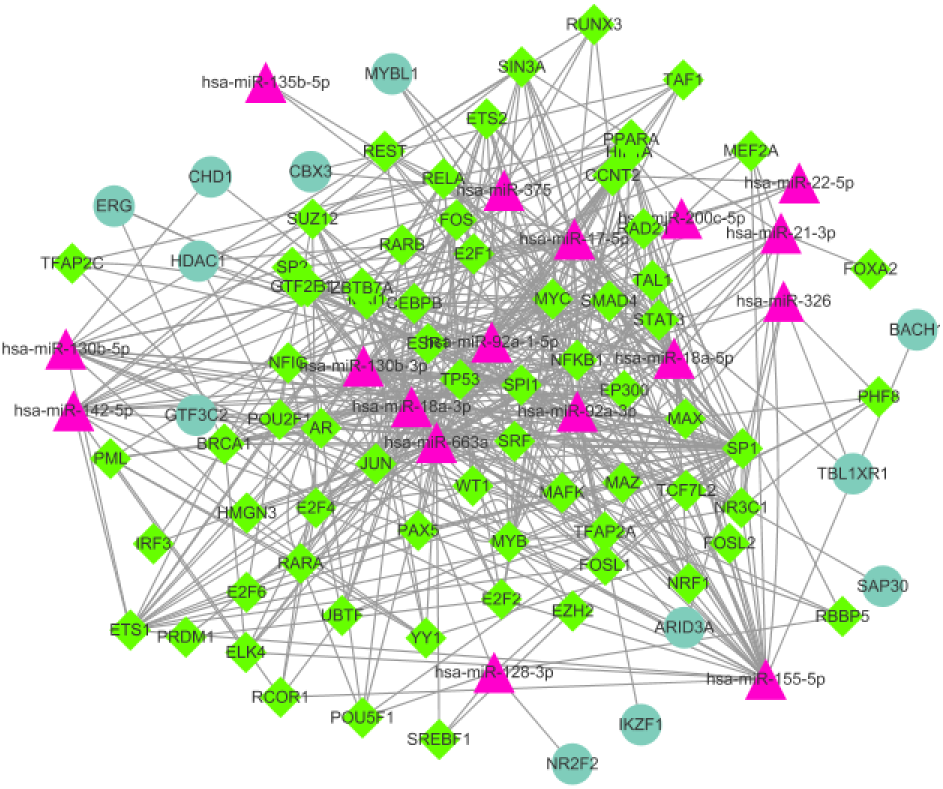
A sub-network of the constructed complex gene regulatory network. The triangles represent the miRNAs, the circles represent genes and the rhombi represent TFs.

### Case study 2: Predicting miRNA targets by integrating different data sources

In this scenario, we would like to predict miRNA targets in Breast cancer with a computational method, e.g. Pearson correlation using expression data from TCGA BRCA together with the target binding information from TargetScan. We then use miRTarBase v7 to validate the predictions of miRNA targets. The obstacle is the inconsistent annotations of miRNAs from different sources. The annotations of miRNAs in TCGA miRNA illumina Hiseq platform are precursors, and the annotations of miRNAs in TargetScan 7.0 and miRTarBase v7 come from different miRBase versions. We apply the *checkMiRNAVersion()* in *miRBaseConverter* to cope with the miRNA name conversion for miRNA expression dataset in TCGA and unify the miRNA version of TargetScan 7.0 and miRTarBase v7 to miRBase v21.

The Level 3 TCGA BRCA RNASeqv2 and miRNAHiseq data are downloaded from the Broad GDAC Firehose using TCGA Assembler [38]. We extract 200 matched tumor samples of gene expression and miRNA expression dataset for our experiments. To reduce computational complexity, we further extract the top 8000 genes based on the big variance among samples in gene expression while keep all the features in the miRNA expression dataset because of the small number of miRNAs. The miRNA precursor names are used to represent each of the miRNAs in the miRNA expression data. We apply *miRBaseConverter* to check the miRBase version number and convert precursor names to the corresponding mature names. The miRNA annotations of TargetScan 7.0 and miRTarBase v7 are based on miRBase version 17 and version 21, respectively. Here we apply *miRBaseConverter* to transform the all miRNA names to the names used in the latest miRBase version 22. So all the experiments data are consistent with the same miRBase version.

For demonstration, we apply the method of Pearson Correlation implemented in the *miRLAB* package to predict the targets of a miRNA base on expression data. The *miRLAB* package provides a complete set of pipelines for predicting miRNA-mRNA regulatory relationships using expression data [20]. To evaluate the performance of a computational method (Pearson correlation in this case), we validate the predicted results using the experimentally confirmed databases for miRNA targets. Therefore, we use miRTarBase v7 to validate the top 100 targets of *hsa-miR-224-5p* predicted by the Pearson correlation method. There are 8 predicted miRNA-mRNAs interactions confirmed in miRTarbase v7, as shown in Table 3.

**Table 3.** The 8 predicted miRNA-mRNAs interactions of *hsa-miR-224-5p* are confirmed in miRTarbase v7

| miRNA | mRNA | Correlation |
| --- | --- | --- |
| hsa-miR-224-5p | BCL2 | -0.45568915 |
|  | FOSB | -0.18087381 |
|  | DIO1 | -0.17142808 |
|  | SLC12A5 | -0.12755148 |
|  | ZKSCAN8 | -0.09462372 |
|  | EYA4 | -0.08509764 |
|  | PTX3 | 0.25557907 |
|  | EFNA3 | 0.22785473 |

Based on the validation result, the correlation between *hsa-miR-224-5p* and some of its confirmed target mRNAs are positive or near zero, while the common knowledge is that miRNAs down-regulate their target mRNAs and hence negative correlation should be expected. The results suggest that the predicted targets by Pearson correlation may include false discoveries. On the other hand, due tothe incompleteness of miRTarbase v7, some targets predicted by the Pearson correlation method (the ones with negative correlation coefficients) are not validated by miRTarBase v7. Therefore, follow-up analyses and/or experiments should be conducted. miRNA regulates gene expression by binding to the target mRNAs. Therefore, the miRNA sequence information is very important for predicting target mRNAs by studying the presence of mRNA conserved sites that match the seed region of each miRNA. In the case that one would like to see the sequence of miRNAs and check with the sequence of predicted mRNAs, the user can simply use the function *getMiRNASequence()* in *miRBaseConverter* for obtaining the multiple sequence of the interest miRNAs. We provide the complete processing scripts in online supplementary data.

## Conclusion

In this paper, we have presented the framework and the usages of the *miRBaseConverter* R package which has been released in the Bioconductor repository. A Shiny-based web application of *miRBaseConverter* has also been presented in the paper for non-R users. The Bioconductor package and the Shiny-based web application provide a suite of functions for converting and retrieving miRNA name, accession, sequence, species, version and family information in different miRBase versions. To our knowledge, there was previously no available tool that could conduct the batch retrieval of miRNA family information, thus representing a challenge in the miRNA-related research. *miRBaseConverter* is a full-scale tool for converting and retrieving information of all miRNAs defined in miRBase. We also have demonstrated the use of the main functions using examples and some case studies. We believe that the package will serve a wide range of applications in miRNA research and provide a full view of the miRNAs of interest.

## Availability and requirements

**Project name**: miRBaseConverter

**Project home page**: http://bioconductor.org/packages/miRBaseConverter/

**Operating system(s)**: Platform independent

**Programming language**: R

**Other requirements**: R (>= 3.4)

**License**: GNU GPL-2

**Any restrictions to use by non-academics**: none

## Acknowledgements

We thank Dr. Martin Morgan, the Bioconductor Project leader, for his valuable comments on the codes to greatly improve the retrieval efficiency of the *miRBaseConverter* package.

## Availability of data and material

The datasets generated and/or analysed during the current study are available in the github depository at https://taoshengxu.github.io/ExperimentData/miRBaseConverter. *miRBaseConverter* is freely available and can be downloaded from Bioconductor depository. *miRBaseConverter* is compatible with Linux, MacOSX and Windows.

## Author’s contributions

TX, TDL and JL conceived the research. TX, NS, JZ and TDL designed and implemented the method and performed the experiments and validation analysis. TX, NS and LL wrote the paper. All authors contributed to the editing of the manuscript. All authors read and approved the final manuscript.

## Ethics approval and consent to participate

Not applicable.

## Consent for publication

Not applicable.

## Additional Files

Additional file 1 — The vignette of miRBaseConverter.

